# Covalent Binding of Boronic Acid-Based Inhibitor GSK4394835A to Phosphodiesterase 3B, a Drug Target for Cardiovascular Disease

**DOI:** 10.1101/2025.10.22.684020

**Authors:** Samuel A. Eaton, David W. Christianson

## Abstract

Boronic acids have become increasingly prominent in the pharmacopoeia ever since the US Food and Drug Administration approved Bortezomib (Velcade) for the treatment of multiple myeloma more than 20 years ago. Since then, four additional boronic acid-based drugs have been approved for clinical use in the treatment of cancer, fungal and bacterial infections, and eczema. The boron atom is inherently electrophilic due to being electron deficient, so the mode of inhibition by a boronic acid very often involves addition of a nucleophile, such as an enzyme-bound residue or a water molecule, to yield a tetrahedral boronate anion. Recently, a new class of boronic acid-based inhibitors of human phosphodiesterase 3B (PDE3B) have been reported, along with the 2.7 Å-resolution crystal structure of PDE3B complexed with one of these inhibitors, GSK4394835A [Rowley et al. (2024) Discovery and SAR study of boronic acid-based selective PDE3B inhibitors from a novel DNA-encoded library. *J. Med. Chem. 67*, 2049–2065]. Surprisingly, the boronic acid moiety of the inhibitor was reported to bind as an intact, unreacted boronic acid. Additional discrepancies were evident in the reported refinement statistics, so we downloaded structure factor amplitudes from the Protein Data Bank (accession code 8SYC) and re-solved the structure. Our initial electron density maps clearly indicated nucleophilic addition of H737 to the boronic acid moiety of the inhibitor, generating a tetrahedral boronate anion in the active site. We re-refined the structure with excellent refinement statistics, concluding that GSK4394835A is a reversible covalent inhibitor of PDE3B.

---

Boron has been an increasingly prominent element in medicinal chemistry ever since the approval of the first boron-containing drug, the peptide boronic acid Bortezomib (Velcade), for the treatment of multiple myeloma in 2003.^1-3^ Bortezomib is a proteasome inhibitor with a unique mode of action: the boronic acid moiety undergoes nucleophilic attack by the hydroxyl group of T1 to yield a covalently-bound tetrahedral boronate anion in both the chymotryptic and caspase-like active sites.^4^ Given that the boronic acid moiety is isosteric with the scissile amide group of an actual peptide substrate, the tetrahedral boronate anion mimics the tetrahedral intermediate and its flanking transition states in the amide hydrolysis reaction (Figure 1).

**Figure 1.**
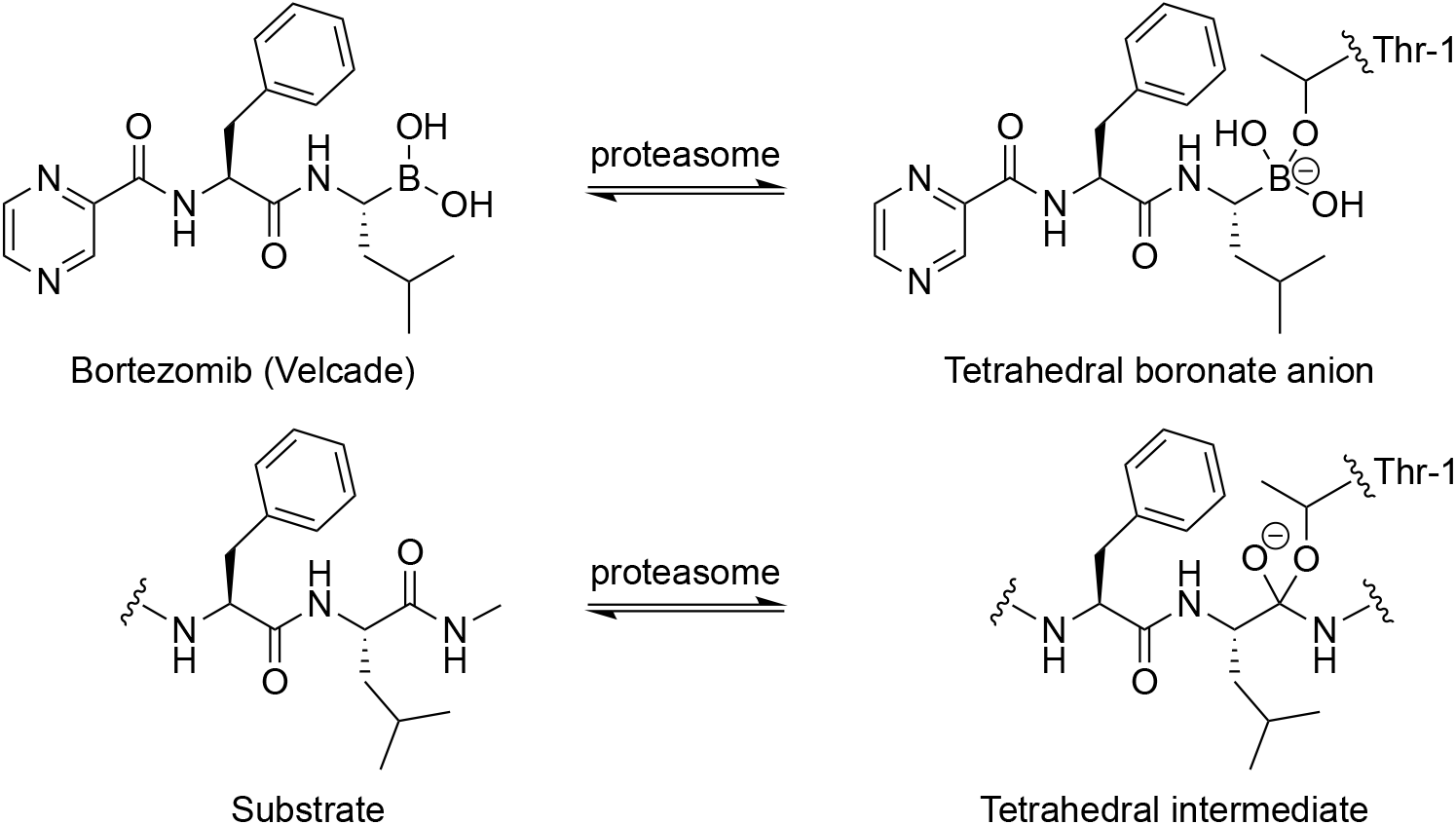
Bortezomib undergoes nucleophilic attack by the hydroxyl group of T1 in the active site of the proteasome to yield a tetrahedral boronate anion. This chemistry mimics that of the first step of catalysis with a hypothetical peptide substrate.

Similar chemistry is observed in the binding of boronic acid inhibitors to other hydrolytic enzymes. For example, boronic acid-based substrate analogues similarly undergo nucleophilic attack by the hydroxyl group of a catalytic serine to yield covalently-bound tetrahedral boronate anions in the active sites of serine proteases and β-lactamases.^5-8^ Such tetrahedral species mimic the tetrahedral transition state in the first step of catalysis by each enzyme. Moreover, such boronic acid chemistry is not limited to hydroxyl nucleophiles: certain boronic acid inhibitors of mandelate racemase undergo nucleophilic attack by an active site histidine residue to yield a covalent adduct.^9^ The reactivity of the boronic acid in these examples is a consequence of the electron-deficient boron atom, which lacks an octet of electrons – nucleophilic addition completes the electron octet around boron.

Cyclic nucleotide phosphodiesterases are metalloenzymes that catalyze the hydrolysis of phosphodiester linkages in cyclic adenosine monophosphate and/or cyclic guanosine monophosphate, both of which are second messengers in myriad cellular pathways. To date, eleven phosphodiesterase isozymes have been identified, several of which are validated drug targets.^10,11^ Of particular interest is human phosphodiesterase 3B (PDE3B), which is a possible drug target for cardiovascular indications.^12-14^ The 2.4 Å-resolution crystal structure of PDE3B was first reported more than 20 years ago and serves as a useful reference point for structural studies of enzyme-inhibitor complexes.^15^

The discovery of novel boronic acid-based inhibitors of PDE3B was recently disclosed along with the 2.7 Å-resolution crystal structure of PDE3B complexed with one of these inhibitors, GSK4394835A (pIC_50_ = 6.2).^16^ Curiously, the structure of the enzyme-inhibitor complex shows an intact, trigonal planar boronic acid bound in the active site (Figure 2).^16^ In view of the inherent reactivity of the electron-deficient boronic acid, it was surprising to see that it did not bind as a tetrahedral adduct with a nearby Lewis base such as H737. The binding of a tetrahedral boronate anion formed with metal-bound water could have been an alternative possibility, as observed for boronic acid binding to the Mn^2+^_2_ cluster in the active site of arginase.^17-19^ It is not unprecedented for an intact boronic acid to bind in an enzyme active site,^9^ but this seems to be more of an exception rather than a rule.

**Figure 2.**
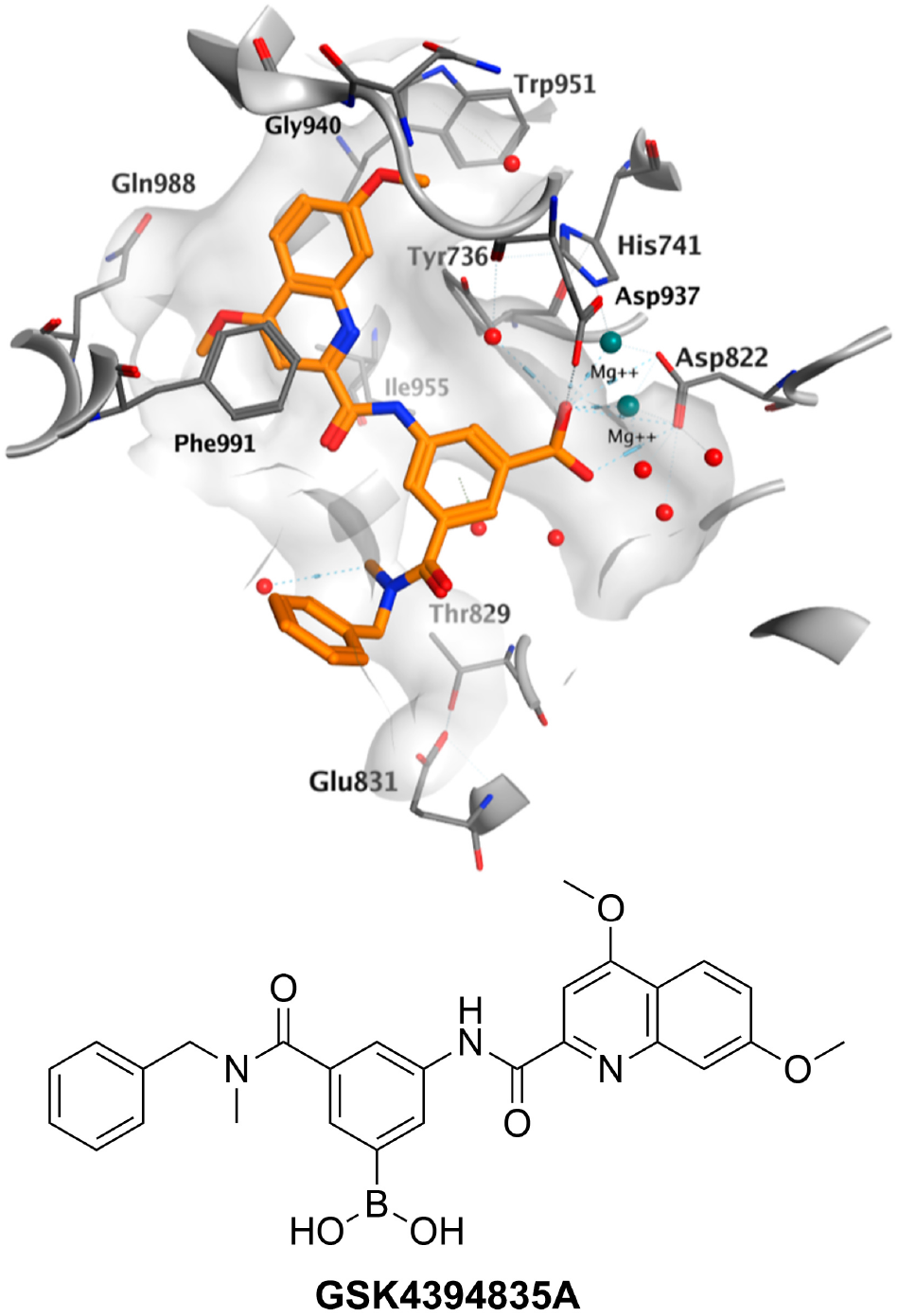
Crystal structure of the PDE3B-GSK4394835A complex (PDB 8SYC). Reprinted from ref. 16, in which the inhibitor is disclosed as compound **23**. The boronic acid moiety is in its trigonal planar form and is shown to interact with each Mg^2+^ ion as well as metal ligands D822 and D937 (note: the inhibitor boron atom is the same orange color as carbon). Copyright 2024 American Chemical Society.

Upon reviewing the data collection and refinement statistics published for the crystal structure determination of the PDE3B-GSK4394835A complex,^16^ we noticed some unusual features, including R_work_/R_free_ = 0.2350/0.2269 and a root-mean-square deviation (rmsd) of 0.068 Å from ideal bond lengths; ordinarily, this value should be less than 0.02 Å. Additionally, the electron density map in the validation report archived with the Protein Data Bank (PDB) reveals extra density around the boron atom of GSK4394835A.^20^ To study the structure of the PDE3B-GSK4394835A complex more closely, we downloaded the atomic coordinates and structure factor amplitudes deposited in the PDB (accession code 8SYC).^20^ Surprisingly, the two coauthors of the PDB deposition^20^ were not included as coauthors of the published paper.^16^ Moreover, many (but not all) of the data collection and refinement statistics recorded in the PDB deposition differ substantially from those reported in the published paper.

In light of our growing concerns regarding the chemistry and crystallography reported for this system,^16^ we re-solved the structure of the PDE3B-GSK4394835A complex by molecular replacement. We used the atomic coordinates of PDE3B (PDB 1SOJ)^15^ as a search probe for rotation and translation function calculations using Phaser^21^ as implemented in PHENIX.^22^ After inspection and adjustment of the initial protein model using Coot,^23^ the structure was refined using PHENIX.refine.^22^ Iterative rounds of manual model adjustment and refinement were performed, with solvent molecules added in the later stages of refinement. From the initial stages of electron density map inspection and structure refinement, it was clear that the electron density corresponding to the bound inhibitor was connected to H737 in monomer A in the asymmetric unit of the tetragonal *P*4_1_ unit cell. Electron density in monomer B was ambiguous, likely due to disorder and/or lower occupancy, so this density was left unmodeled. However, this density was similarly connected to H737 in difference electron density maps in which H737 was omitted from the structure factor calculation (Figure S1), suggesting that covalent binding occurs even at low occupancy binding. The atomic coordinates of GSK4394835A were added to monomer A in the final stages of refinement. The structure refined smoothly to convergence with R_work_/R_free_ = 0.190/0.238 and rmsd = 0.002 Å from ideal bond lengths. MolProbity^24^ was employed for structure validation, and final refinement statistics are recorded in Table 1 and compared with statistics listed for the previously-reported structure determination.^16^

**Table 1.** Refinement statistics for the PDE3B-GSK4394835A complex.

| Structure | Ref. 16 | This study |
| --- | --- | --- |
| Reflections used in refinement/test set | 26,785 (2410)/1419 (141) | 18,141 (312)/840 (12) |
| $R_{work}^{a,b}$ | 0.235 (0.392) | 0.190 (0.306) |
| $R_{free}^{a,b}$ | 0.227 (0.375) | 0.238 (0.358) |
| no. of protein chains | 2 | 2 |
| no. of nonhydrogen atoms | 5,876 | 5,527 |
| protein | 5,716 | 5,431 |
| ligand | 78 | 51 |
| solvent | 82 | 45 |
| average *B* factor (Å<sup>2</sup>) | 54 | 49 |
| protein | 54 | 49 |
| ligand | 77 | 40 |
| solvent | 34 | 35 |
| Root-mean-square deviation from ideal geometry |  |  |
| bonds (Å) | 0.068 | 0.002 |
| angles (deg) | 2.51 | 0.5 |
| Ramachandran plot<sup>c</sup> |  |  |
| Favored (%) | 94.13 | 97.41 |
| Allowed (%) | 4.73 | 2.59 |
| Outliers (%) | 1.15 | 0.00 |
| MolProbity score | 1.61 | 1.33 |

<sup>a</sup>Values in parentheses refer to data in the highest shell. <sup>b</sup> $R_{work} = \sum ||F_o| - |F_c|| / \sum |F_o|$ for reflections contained in the working set. $|F_o|$ and $|F_c|$ are the observed and calculated structure factor amplitudes, respectively. $R_{free}$ is calculated using the same expression for reflections contained in the test set held aside during refinement. <sup>c</sup>Calculated with MolProbity; although not reported for PDB 8SYC in ref. 16, atomic coordinates were subject to Molprobity analysis and the resulting score reported here.

The structures of monomers A and B of the PDE3B-GSK4394835A complex in the asymmetric unit are essentially identical, with the exception of well-ordered inhibitor density in the active site of monomer A but ambiguous inhibitor density in monomer B that was left unmodeled. The rmsd = 0.224 Å for 315 Cα atoms between monomers A and B. Importantly, the re-solved and re-refined structure of the complex clarifies the binding mode of the boronic acid moiety in monomer A: it does not bind as a trigonal planar boronic acid as initially presented,^16^ but instead binds as a covalent adduct with H737 to form a tetrahedral boronate anion (Figure 3). Therefore, GSK4394835A is a covalent inhibitor of PDE3B and it is reversible in view of the inherent reversibility of nucleophilic addition to boron. The authors report that they “were unable to engineer away the boronic acid group, without compromising compound potency”.^16^ The covalent bond between the boronic acid moiety of GSK4394835A and H737 is the likely reason why.

**Figure 3.**
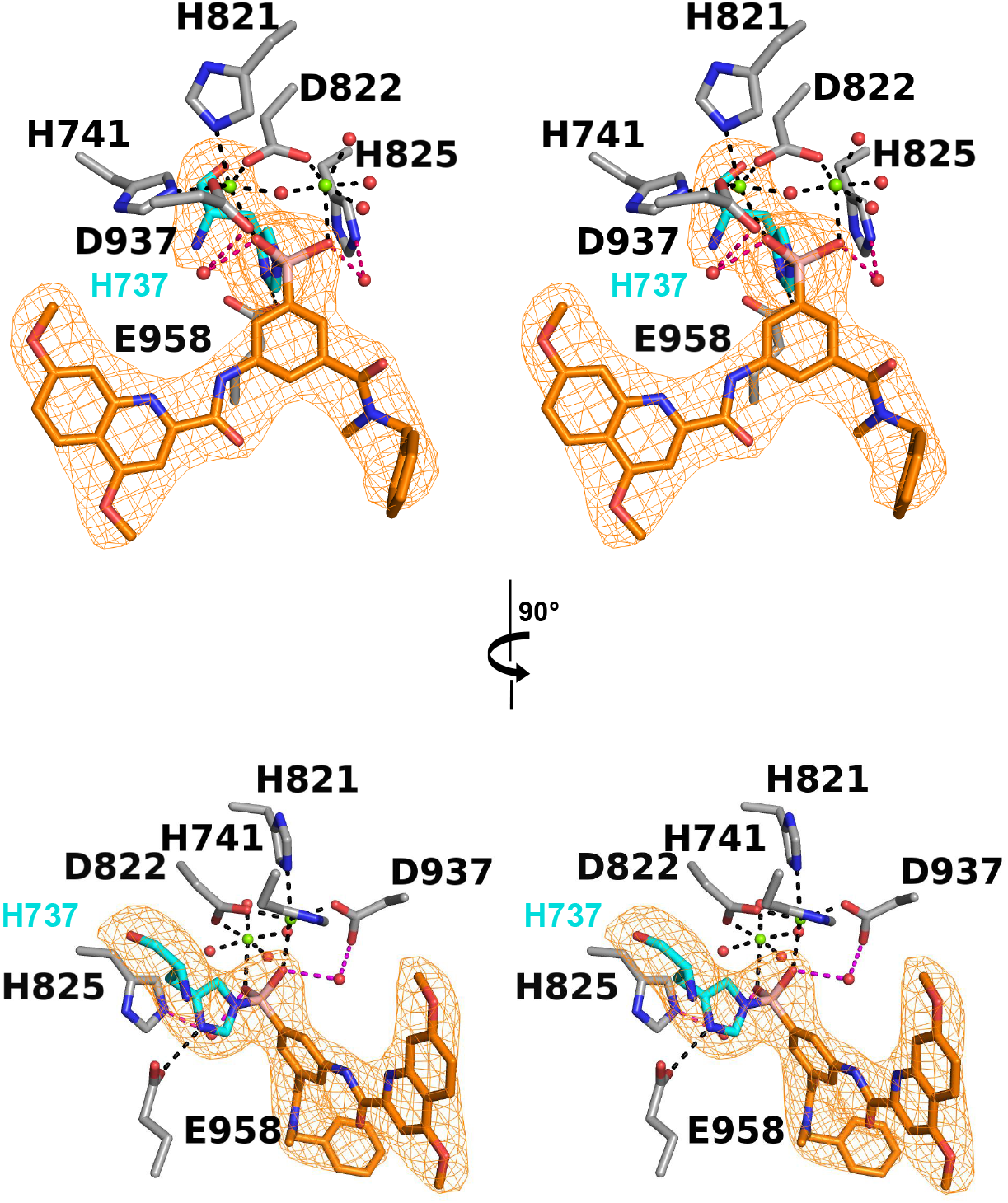
Two orthogonal stereoviews of the PDE3B-GSK4394835A complex (PDB 9YUD) showing a Polder omit map of the covalent adduct between H737 and GSK4394835A, contoured at 4.0 σ. The electron density between H737 and the boronic acid moiety is continuous and strong, indicative of a covalent linkage.

In addition to the zwitterionic Nε^+^–B^−^ bond formed between H737 and GSK4394835A, there are several intermolecular interactions of interest in the enzyme-inhibitor complex. Most notably, each boronate hydroxyl group of the inhibitor coordinates to a Mg^2+^ ion. The octahedral coordination sphere of Mg_A_^2+^ is completed by H741, H821, D822, and D937, plus a metal-bridging solvent molecule. The octahedral coordination sphere of Mg_B_^2+^ is completed by the metal-bridging solvent molecule, D822, and three water molecules (Figure 4). Additionally, water molecules #48 and #50 make bridging hydrogen bond interactions between boronate hydroxyl groups and the sidechains of D937 and H825, respectively. No other enzyme-inhibitor hydrogen bond interactions are observed, but the dimethoxyquinoline moiety of the inhibitor engages in an offset π-stacking interaction with the sidechain of F991.

**Figure 4.**
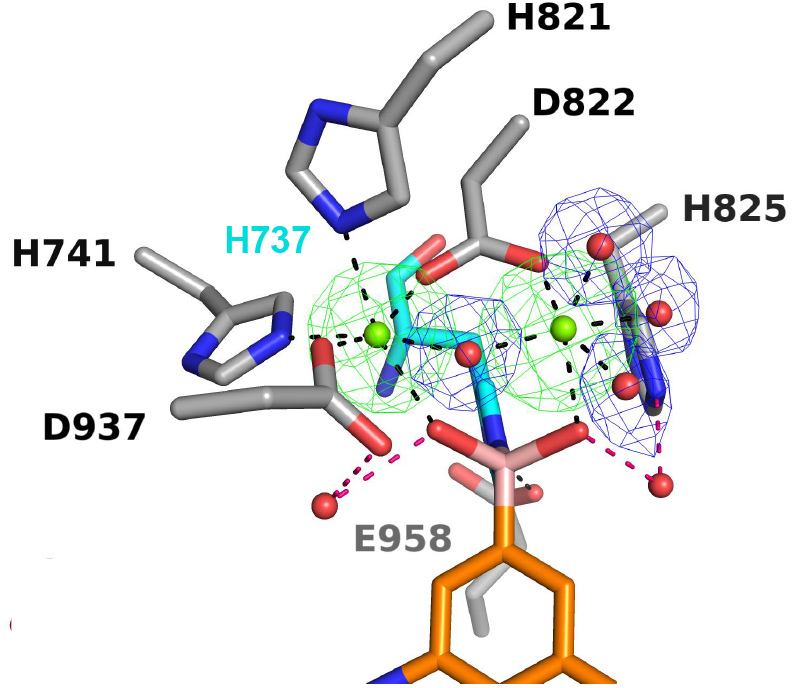
Polder omit maps of the Mg^2+^ ions and metal-bound solvent molecules contoured at 5.5 σ and 5.0 σ, respectively, in the PDE3B-GSK4394835A complex (PDB 9YUD). Metal coordination geometries are distorted octahedral. Metal coordination and hydrogen bond interactions are shown as black and magenta dashed lines, respectively.

Since H737 donates a hydrogen bond to E958, the basicity (and nucleophilicity) of the H737 imidazole group is enhanced. For example, a carboxylate-histidine hydrogen bond elevates the pKa of the histidine imidazole by approximately 1.5 log units.^25,26^ The formation of a covalent bond with H737 is a novel and unanticipated mode of action for GSK4394835A.

In conclusion, reanalysis of the X-ray crystallographic data deposited in the PDB for the PDE3B-GSK4394835A complex^16^ indicates that the mode of inhibition is covalent and reversible. Key binding interactions of the boronic acid pharmacophore are summarized in Figure 5. In view of the unanticipated nucleophilicity of H737, it is possible that other boronic acid derivatives,^16^ as well as other reactive pharmacophores such as ketones and aldehydes, will similarly bind in reversible covalent fashion. The newly discovered chemistry of boronic acid binding to PDE3B may point to exciting new inhibitors designs in the continuing exploration of potential new therapeutics for the treatment of cardiovascular disease.

**Figure 5.**
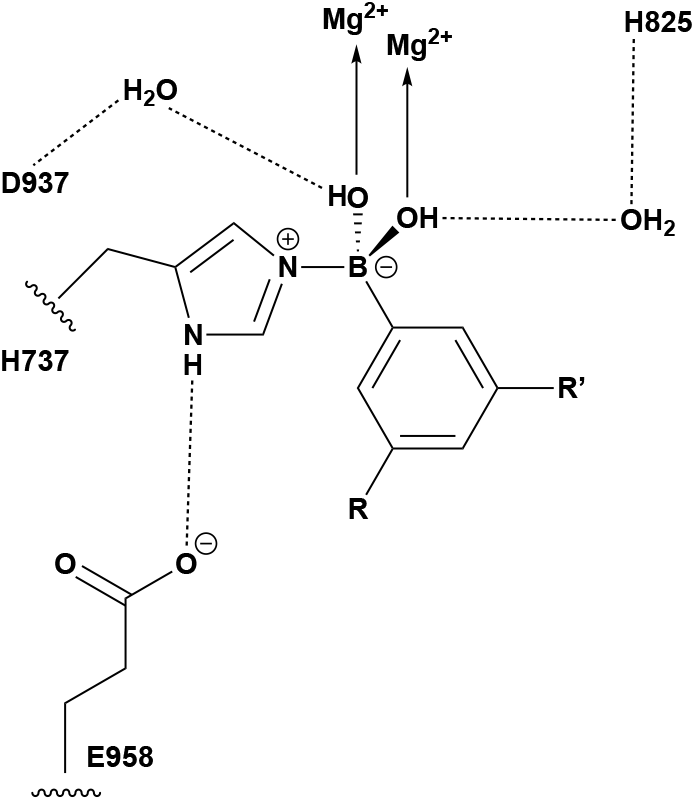
Key binding interactions of the boronic acid pharmacophore of GSK4394835A. Metal coordination and hydrogen bond interactions are indicated by arrows and dashed lines, respectively.

## ASSOCIATED CONTENT

### Supporting Information

The Supporting Information is available free of charge on the ACS Publications website at DOI: Figure S1, polder omit electron density map of H737 in the active site of monomer B.

### Accession Code

The atomic coordinates of the PDE3B-GSK4394835A complex have been deposited in the PDB with accession code 9YUD and will be released upon publication.

## ACKNOWLEDGEMENTS

D.W.C. is grateful to his Ph.D. adviser, the late Professor William N. Lipscomb, Jr. at Harvard University, for his lasting mentorship in protein crystallography as well as boron chemistry.

## For TOC use only

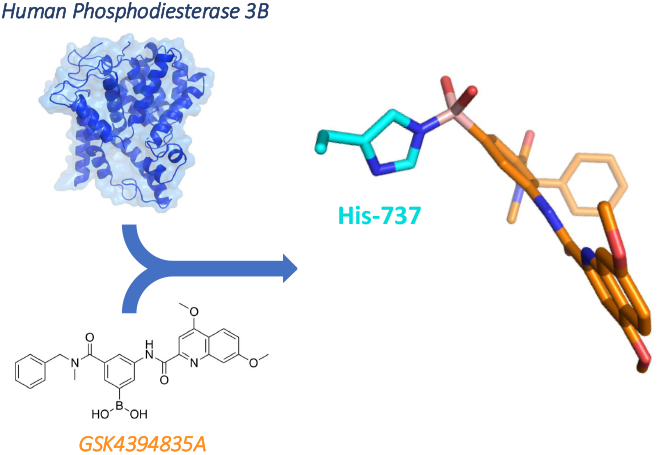

