## Supporting Information for "Covalent Binding of Boronic Acid-Based Inhibitor GSK4394835A to Phosphodiesterase 3B, a Drug Target for Cardiovascular Disease"

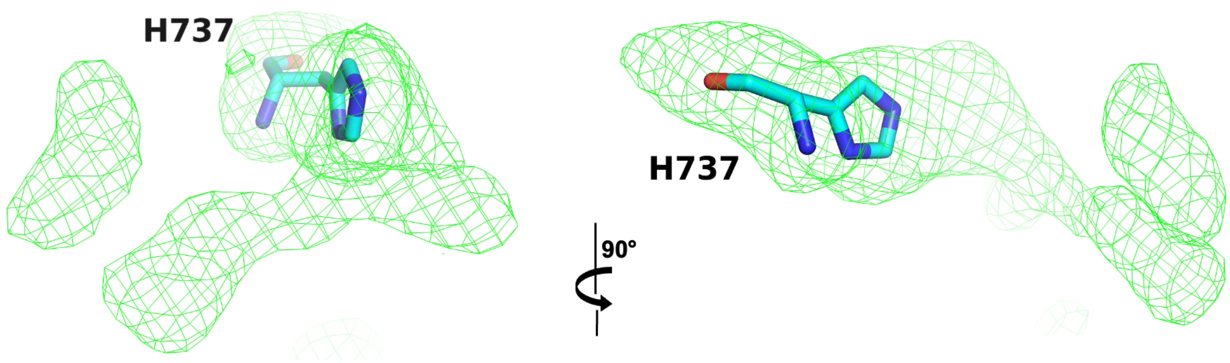


**Figure S1.** Polder omit map of H737 in monomer B contoured at 3.0 σ. Electron density for GSK4394835A is weak and discontinuous, perhaps due to disorder or partial occupancy. Accordingly, the inhibitor was not built into this density. However, the continuous density between H737 and the inhibitor suggests that the inhibitor binds covalently in the active site of monomer B, just as it does with higher occupancy in monomer A.
